# Deficiency of miR130a leads to fat hypertrophy, hepatic steatosis, insulin resistance and glucose intolerance in mice

**DOI:** 10.64898/2026.01.24.701468

**Authors:** Yi-Cheng Chang, Ching-Han Chuang, Shu-Fan Chou, Jyun-Yuan Huang, Chia-ho Shih

**Affiliations:** Institute of Biomedical Science, Academia Sinica, Taipei, 115 Taiwan; Department of Internal Medicine, National Taiwan University Hospital, Taipei, 100, Taiwan; Graduate Institute of Medical Genomics and Proteomics, National Taiwan University, Taipei, 100, Taiwan; Graduate Institute of Microbiology, College of Medicine, National Taiwan University, Taipei, 110, Taiwan; Graduate Institute of Medicine, Kaohsiung Medical University, Kaohsiung, 807, Taiwan; Graduate Institute of Cell Biology, China Medical University, Taichung, 406, Taiwan

## Abstract

Insulin resistance, excessive and ectopic fat accumulation, chronic low-grade inflammation, and pancreatic beta-cell failure are pathological features of type 2 diabetes mellitus.MiR-130a has been demonstrated to suppress the mRNA levels of PPARγ, NF-κB, and TNF-α *in vitro*. PPARγ is a master regulator of systemic fat and glucose metabolism. NF-κB and TNF-α are pivotal modulators of inflammation. Therefore, we aimed to examine the systemic effect of miR130a on fat metabolism, glucose/insulin homeostasis, and inflammation in mice.

We found that mirR130a-deficient mice exhibited larger white fat mass with hypertrophic adipocytes, increased lipogenic gene expression in fat, and elevated serum leptin levels than controls. The white fat pads of mirR130a-deficient mice showed significant macrophage infiltration with enhanced expression of pro-inflammatory genes. In addition, mirR130a-deficient mice had more severe hepatic steatosis and higher hepatic triglycerides content than controls. Similarly, mirR130a-deficient mice had increased macrophage infiltration and lipogenic and inflammatory gene expression in the liver. Consistently, we found that *Lep*^ob/ob^ mice expressed markedly decreased miR130a expression in the liver and white fat compared to controls.

Importantly, mirR130a-deficient mice displayed impaired glucose tolerance and worsened insulin resistance, accompanied with reduced serum adiponectin levels. Furthermore, insulin secretion is reduced in mirR130a-deficient mice compared to controls.

In conclusion, knockout of miR130a in mice results in fat hypertrophy, hepatic steatosis, increased macrophage infiltration in liver and fat, glucose intolerance, and insulin resistance. These data indicate miR130a exert systemic anti-diabetic effects.

## Introduction

Type 2 diabetes mellitus is a chronic progressive disease characterized by insulin resistance and beta-cell failure. Insulin resistance is the fundamental feature of type 2 diabetes. Two major theories-the “lipid spillover” and the “inflammation” theories are widely accepted as the underlying mechanism of insulin resistance (1,26). The “lipid spillover” theory holds that excessive calorie intake leads to excessive fat accumulation in adipose tissue and spillover of excessive fat to ectopic tissue such as liver and muscle. These lipid intermediate metabolites such as ceramides block insulin signaling cascade in liver and muscle, leading to insulin resistance in these tissues (1,2). The “inflammation” theory is based on the observation that fat expansion leads to fat cell necrosis and macrophage infiltration into fat tissues, triggering low-grade systemic inflammation and the release of pro-inflammatory cytokines from fat tissue into circulation. These pro-inflammatory cytokines further activate stress kinases that impair insulin signaling in other tissues (1, 2).

Peroxisome proliferator-activated receptor γ (PPARγ) is a master transcriptional regulator of adipogenesis, lipid metabolism, glucose and insulin homeostasis, and adipocytokine release (3). The synthetic PPARγ ligands thiazolidinediones are currently important anti-diabetic drugs. However, little is known about the regulation of PPARγ expression. MicroRNAs are highly-conserved, small non-coding, single-strand RNA of ∼ 22 nucleotides that typically bind and degrade target mRNA or inhibit the translation of target mRNA by complementary sequences. Through fine-tuning of expression of multiple target mRNAs, microRNAs are involved in various physiological and pathological processes (4). Approximately 2000 miRNAs have been identified across the human genome with ∼ 45000 miRNA target sites within human 3’UTRs (5). We and other groups have previously demonstrated that miR130a reduced PPARγ expression, probably via binding to its coding region and 3’UTR (6–11). Inhibition of miR130a increased PPARγ expression, promoted adipocyte differentiation, and enhanced down-stream adipogenesis gene expression while miR130a over-expression decreased PPARγ down-stream genes including adiponectin and leptin (7). Another study showed that knockdown of miR130a decreased glucose-stimulated insulin secretion in pancreatic beta-cell lines *in vitro* (12). Furthermore, several studies showed that miR130a also repress NF-κB/p65 (13, 14) and TNF-α mRNA expression (15, 16, 17, 18). These data suggest miR130a may regulate systemic fat and glucose metabolism and pro-inflammatory responses.

In current study, we aimed to explore the systemic effect of miR130a on fat metabolism and glucose/insulin homeostasis using miR130a-deficient mice. We found that miR130a-deficient mice exhibited fat hypertrophy, hepatic steatosis, glucose intolerance, insulin resistance, and increased inflammation in fat and liver. These data indicate miR130a is an important regulator for systemic fat metabolism and glucose homeostasis.

## Methods

### Transcription activator-like effector nucleases (TALEN) target site selection and TALEN construction

To target the seed region of the mouse miR-130a locus, various TALEN pairs were designed using an online tool, TAL Effector Nucleotide Targeter 2.0 (19). To design efficient miRNA targeted TALENs, we followed these criteria: (1) TALEN binding sites were set to 20 bp, including the first T, to ensure high specificity of gene targeting; (2) spacer lengths of 13–20 bp were chosen to maximize cleavage efficiency; (3) the miRNA seed sequence was situated centrally within the spacer to direct cleavage to the seed region (20). Following these criteria, the target sequences of TALENs for mmu-mir-130a are as follows; L1: 5’-TGTGCTACTGTCTAACGTG-3’, L2: 5’-TGCTACTGTCTAACGTGTA-3’ and L3: 5’-TACTGTCTAACGTGTACCG-3’; right arm: 5’-TGCCCTTTTAACAT-3’ and 5’-TACAAGGCCGATGCCC-3. We used custom TALEN service (Cold Spring biotech Co.Ltd) to construct plasmids coding for TALEN with an EF1-driven expression cassette using the SIDANSAI TALEN Assembly Kit (Cat. No: 1801 −060, 1802-030, and 1803-015; SIDANSAI Biotechnology Co. Inc). The final constructs were confirmed by DNA sequencing.

### Evaluation of TALENs activities in cultured cell line

Briefly, a total of 4 × 10^5^ Hepa 1-6 hepatoma cells were transfected with 0.5 μg of each TALEN-encoding plasmid and cultured for 24 hrs. The puromycin was then added to the culture medium (final concentration 2 μg/mL), to kill untransfected cells. After 4 days of puromycin selection, puromycin-resistant clones were expanded for further DNA sequencing and RNA analysis.

### Generation of *miR130a*-deficient mice by microinjection

TALEN plasmids were linearized by Not1and HindIII endonuclease digestion respectively. The procedures of linearized plasmid used as a template for *in vitro* transcription reaction and microinjection were manipulated by Institute of Molecular Biology Academia Sinica transgenic mouse core facility.

### Mouse genotyping

Tail genomic DNA of TALEN-transfected mice was isolated with REDxtract-N-Amp Tissue PCR Kit (Sigma Aldrich, St. Louis, MO, USA), done according to the manual. Genotyping miR-130a heterozygous and homozygous mutant mice were identified by performing PCR using following primers: mmu-miR-130aKO forward: 5’-TCTTTCTCCTGCCTAAGCACCT-3’; mmu-miR-130a KO reverse: 5’-GGGAGGGCTCCATATATCCAAAT-3’. The PCR program consists of a denaturation step at 95°C for 4 min, followed by 34 cycles of denaturation (94°C for 30 s), annealing (55°C for 30 s) and extension steps (72°C for 60 s). The program ends with a completion step at 72°C for 180 s. Each PCR tube contains 0.05 U of MyTaq™ HS DNA Polymerase in 14μL of reaction mix (react ion buffer, dNTPs and primers) and 100 ng of genomic DNA in final volume of 15 μL. The PCR products were purified by PCR clean-up kit for sequencing.

### Animal protocols, housing, and diets

C57BL/6J (B6/J) and C57BL/6J-miR-130a ^−/−^ (B6/J-miR-130a ^−/−^) were bred in the animal facility of the Institute of Biomedical Sciences, Academia Sinica. B6/J-miR-130a ^−/−^ mice were generated in our laboratory and backcrossed to B6/J mice for at least five generations.All animal experiments were performed according to institutional ethical guidelines and were approved by the Institutional Animal Care and Use Committee (IACUC) of the Academia Sinica, Taiwan (IACUC number: 16-12-1017). All mice were housed under standard condition at 23°C and 12/12 hr light/dark cycle in animal centers of Institute of Biomedical Science (IBMS) of Academia Sinica. Mice were fed on high-fat-high-sucrose diet (Envigo TD.88137) or chow diet (PicoLab® Rodent Diet #205053).

### Glucose and insulin tolerance test

Glucose tolerance was evaluated by oral (OGTT) and intraperitoneal glucose tolerance test (ipGTT) after a 6-hour fasting at the age of 20 weeks. For OGTT, glucose water (1mg/g) was given by oral gavage and tail blood glucose was measured with a glucometer (Contour Plus, Bayer) at 0, 15, 30, 45, 60, 90, 120 min. For ipGTT, tail blood glucose was measured at 0, 15, 30, 45, 60, 90, 120 min after intraperitoneal injection of glucose water (1mg/kg). For insulin tolerance test (ITT), mice were fasted 4 hr and then intraperitoneal injected with 0.8 IU/kg of insulin (Humulin R, Eli Lilly) at the age of 20 weeks. Tail blood glucose was measured at 0, 15, 30, 45, 60, 90, 120, and 180 min.

### Oil-Red O stain

The mouse liver was fixed in10% neutral formalin overnight then rehydrated in 30% sucrose overnight. The liver was then frozen in OCT gel and two sections were obtained for each mouse. The stock of Oil-Red O (Sigma #O0625) was prepared as 0.5% (wt/vol) in isopropanol. Precipitates were removed by filtration with Whatman paper. Before staining, the working solution was prepared by mixing 60 % stock with 40 % H_2_O (vol). Liver section were washed with phosphate-buffered saline (PBS), fixed with 3.7% of formaldehyde for 2 min, and then washed with PBS again. Oil-Red-O working solution was added for 10 min and then aspirated. Liver section were immediately washed with H_2_O and were dried in ambient air. To quantify lipid amount, cells were dissolved with 100% isopropanol and the absorbance was determined at 520 nm.

### Protein extraction and western blotting

Total protein from cells was extracted with RIPA buffer containing 50 mM Tris–HCl, pH 7.4, 150 mM NaCl, 2 mM ethylenediaminetetraacetic acid (EDTA), 50 mMNaF, 1% Nonidet P-40, 0.5% sodium deoxycholate, 0.1% sodium dodecyl sulfate (SDS) and 1 mMphenylmethylsulfonyl fluoride (PMSF) for 5 min, sonicated, and centrifuged to remove debris. For reducing SDS-polyacrylamide gel electrophoresis, the protein sample was mixed with SDS sample buffer containing dithiothreitol (DTT) and denatured by boiling for 5 min. For non-reducing SDS-polyacrylamide gel electrophoresis, the protein sample was reconstituted in non-reducing SDS sample buffer without DTT. Use primary antibodies targeted PPARγ (#sc-7196 Santa Cruze), α-tubulin (#GTX-628802, GeneTex) and secondary antibodies (anti-rabbit-HRP; GeneTex) to probe the proteins then stained with ECL to detect the protein levels of the livers.

### Histology, adipocyte size measurement, and adipocyte number estimation

For hematoxylin and eosin (HE) staining, white adipose tissues was fixed in 10% neutral formalin, processed and embedded in paraffin before sectioning and staining. Two sections were obtained for each mouse. The stained sections were scanned and analyzed using Pannoramic Viewer (https://www.3dhistech.com/pannoramic_viewer) and Image J software (http://rsbwed.nih.gov/ij/). For adipocyte size measurements, 200 consecutive fat cells of the gonadal fat pad from each mouse were selected for the area measurement. The adipocyte number of the gonadal fat pad was calculated as the fat pad volume divided by the average fat cell volume. Fat pad volume was calculated as fat pad weight (g) divided by fat density (0.915g/cm^3^).

### Measurement of insulin signaling

To evaluate insulin signaling *in vivo*, insulin were injected intraperitoneally 15 min before mice were sacrificed. Tissue samples (perigonadal fat, quadriceps muscle, and liver) were extracted with RIPA buffer [50 mM Tris-HCl, pH 7.4, 2 mM ethylenediaminetetraacetic acid (EDTA), 150 mM NaCl, 50 mM NaF, 1% Nonidet P-40, 1 mM phenylmethylsulfonyl fluoride (PMSF), 0.5% sodium deoxycholate, and 0.1% sodium dodecyl sulfate (SDS)] containing phosphatase inhibitor cocktail (cat. no. 04693132001, Roche), and homogenized. Samples were then centrifuged at 13,000 rpm for 10 min at 4°C to remove debris. Samples were separated by SDS - polyacrylamide gel electrophoresis (PAGE), transferred to polyvinylidene difluoride (PVDF) membrane, and probed with anti-phospo-Akt antibody (cat. no. 4058, Cell Signaling) and Akt antibody (cat. no. 9272, Cell Signaling) with secondary anti-rabbit antibody.

### RNA extraction, cDNA synthesis and RT-qPCR

Briefly, tissue was homogenized and extracted by TRIzol reagent (Invitrogen) and 2 μ g of total RNA was reverse transcribed into cDNA using random primers and High Capacity cDNA Reverse Transcription kit (Applied Biosystem, Grand Island, NY) at 37℃ for 120 minutes. The cDNA product was then diluted 100 times for real-time PCR analysis using Power SYBR Green PCR master mix (Applied Biosystem, Grand Island, NY), and the default condition in a 20 μl reaction volume by Applied Biosystems 7500 Real-Time PCR System. Data were analyzed by relative quantification methods (ΔΔCt methods) using 7500 software V2.0.1.Mouse Glyceraldehyde 3-phosphate dehydrogenase (*Gapdh*) mRNA was used as the internal control. Primer sequences are listed in Table 2. All qPCR reactions were run in duplicates for each sample. The primers used are listed in **Supplementary Table 1**

### MicroRNA extraction and stem-loop qPCR

For determination of miRNA expression levels, TaqMan RT and stem-loop real-time assay (Applied Biosystems) were used. Briefly, 100 ng RNAs were reverse transcribed by specific stem-loop primer, miR-130a (assayID: 000454), and further analyzed by Taq man real-time PCR assay using default setting. U6 snRNA (assayID: 001973) was used as an endogenous control. Data were analyzed by Applied Biosystems 7500 software V2.0.1

### Indirect calorimetry

Mice were measured at the age of 7 weeks using an 8-chamber LabMaster Calorimetry Module (TSE-Systems GmbH) with one mouse per chamber. After acclimatization individually for 72 hrs, the O_2_ consumption (VO_2_, mL/kg/min), CO_2_ production (VCO_2_, mL/kg/min), and respiratory quotient (ratio of VCO_2_/VO_2_) were determined. VO_2_ and VCO_2_ were recorded every 30 min for a total of 48 hrs. The energy expenditure was calculated as the product of the calorific value of (3.815 + 1.232×respiratory exchange ratio) and VO_2_. To measure the food intake, mice were housed in metabolic cages for 72 hours, and food and water intake were measured during the final 48 hrs.

### Serum metabolic parameters

Retro-orbital blood was collected after fasting for 6 hr in mice at the age of 24 weeks under anesthesia by inhalation of isofluorane in oxygen. Fasting plasma adiponectin (ab108785, Abcam), leptin (#27160, IBL), MCP-1 (MJE00B, R&D), insulin (#10-1247-01, Mercodia), and TNF-α (MTA00B, R&D) concentrations were measured using ELISA kits.

### Immunohistochemical (IHC) staining

Liver and white adipose tissues were fixed in 10% neutral formalin and then embed into wax to paste the section onto the slide. The slide was probed with antii-F4/80 antibody (Biolegend #123102) and secondary antibodies (anti-mouse-HRP; Dako #K4000) to detect the F4/80 proteins and then DAB staining. Finally, use the hematoxylin to stain the nucleus. The stained sections were scanned and analyzed using Pannoramic Viewer (https://www.3dhistech.com/pannoramic_viewer) and Image J software (http://rsbwed.nih.gov/ij/).

### miRNA Northern blot

To check the miR-130a expression in miR-130a^−/−^ mice, total cellular RNAs including miRNAs were extracted and run the denaturing acrylamide gel and transferred to nylon membrane. The membrane was then hybridized with the 32P labeled DNA oligonucleotide probe which is complementary to the miR-130a-3p and detected with X-ray film.

### Statistical analyses

Independent Student *t*-tests were used for comparisons between two independent groups. Statistical analyses were conducted using GraphPad PRISM 5.0 and SAS 8.0. All two-sided *P*-values < 0.05 were considered as statistically significant.

## Results

To explore the systemic effect of miR130a on fat metabolism and glucose homeostasis, we generated miR130a-deficient mice (miR130a ^−/−^) using the TALEN techniques (**Figure 1a-1c**). As shown in **Figure 1a-c**, the R2/L3 TALEN pair effectively reduced miR130a expression in Hepa 1-6 cells. Therefore, R2/L3 TALEN pair-encoding plasmid was used for generating miR130a-mutant mice (**Figure 1d**). Northern blots showed that mature miR130a mRNA levels were drastically reduced in the liver of miR130a-mutant mice (**Figure 2a**) and stem-loop reverse transcriptase quantitative PCR (RT-qPCR) demonstrated that miR130a level are nearly absent in the liver (**Figure 2b**) as well as other tissues including white fat, brown adipose tissue, skeletal muscle, cerebellum, cerebrum, and the heart of miR130a-mutant mice (**Figure 2c**). In line with our previous results in cell model, PPARγ level was increased in the liver of miR130a-deficient mice measured by immunoblots (**Figure 2d**) and RT-qPCR (**Figure 2e**).

**Figure 1.**
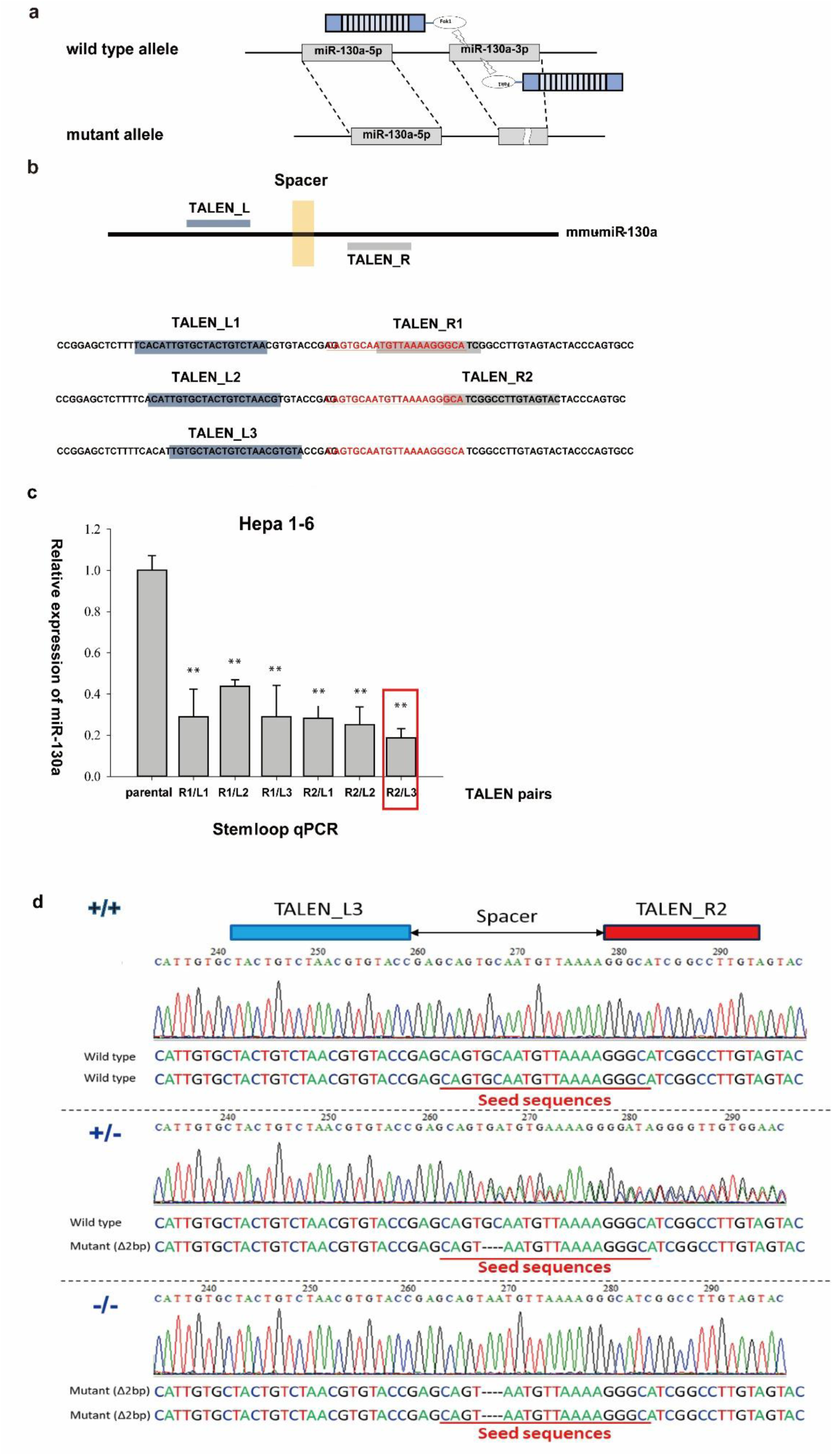
Establishment of miR-130a knockout mice by TALEN approach. (a) The schematic view of the TALEN pair binding on miR-130a −3p site. (b) The illustrations of the five different designed TALEN sequences on mouse miR-130a-3p site. (c) Efficacy of the six TALEN pairs to reduce the miR-130a expression in Hepa 1-6 hepatoma cell lines. (d) PCR genotyping of wild-type (+/+), miR-130a heterozygous knockout mice (+/-) and miR-130a homozygous knockout (−/−) mice. (Δ2bp: two base pair deletions in mmu-miR-130a-3p seed sequences)

**Figure 2.**
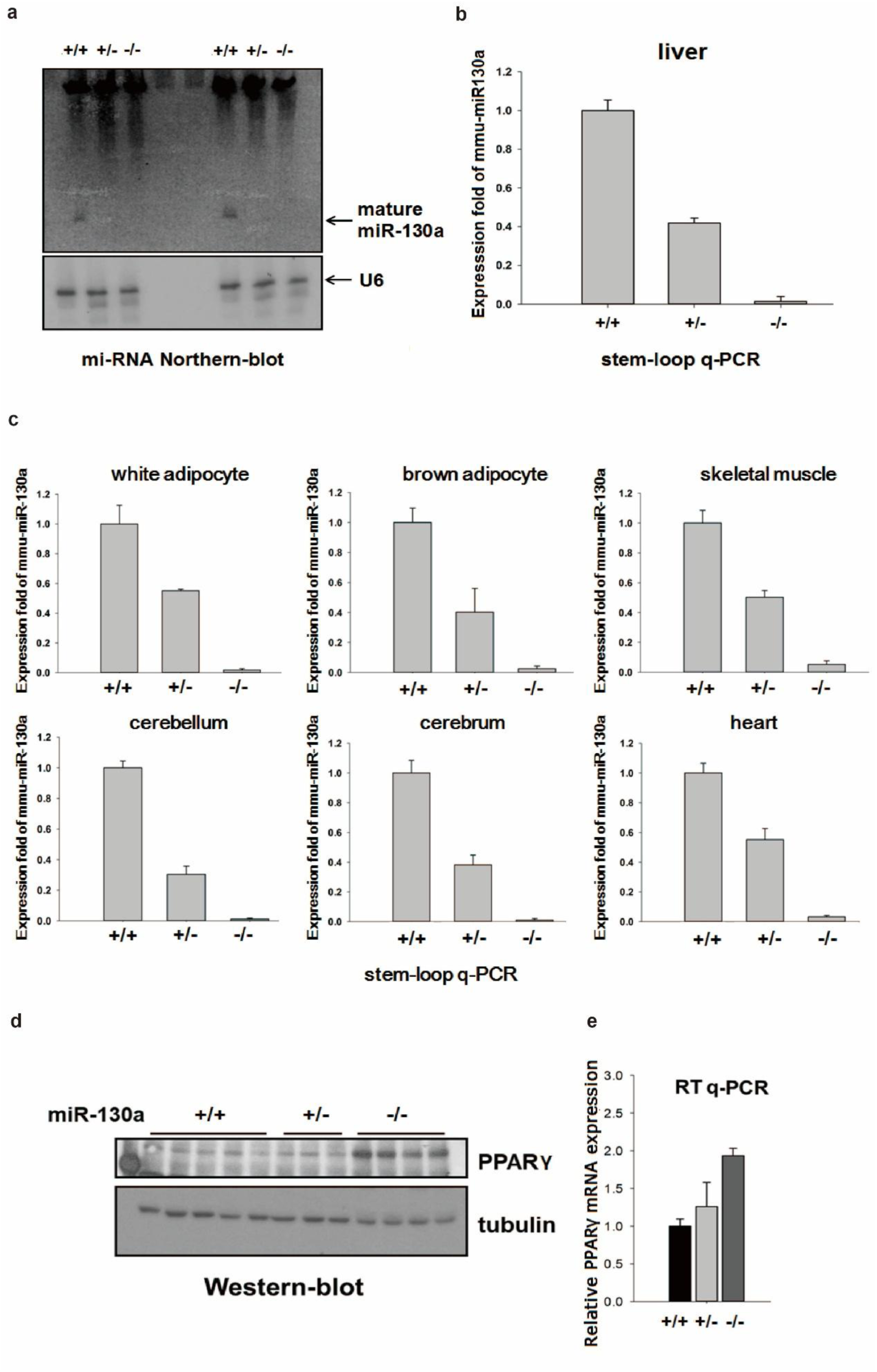
The expression of miR-130a in miR-130aknockout mice. (a) miRNA northern blot and (b) stem-loop qPCR verifying the deletion of miR-130a in the liver of wild-type mice (+/+), miR-130a heterozygous (+/-) and miR-130a homozygous (−/−) knockout mice (c) The expression of miR-130a in white adipose tissue, brown adipose tissue, skeletal muscle, cerebellum, cerebrum, and heart of wild-type mice (+/+), miR-130a heterozygous (+/-) and miR-130a homozygous (−/−) knockout mice (d) PPARγ protein and (e) mRNA levels are up-regulated in miR-130a knockout mice.

There was no difference of body weight between miR130a-deficient mice and wild-type littermates on either high-fat high-sucrose diet (HFHSD) (**Supplementary Figure 1a**) or chow diet (data not shown). There was also no significant difference in energy expenditure (**Supplementary Figure 1b**), respiratory exchange ratio (RER) (**Supplementary Figure 1c**), physical activity measured by running wheels (**Supplementary Figure 1d**), food intake (**Supplementary Figure 1e**) and water intake (**Supplementary Figure 1f**) between mutant and control mice on HFHSD.

However, when fed on HFHSD, miR130a-deficient exhibited significantly larger perigonadal and inguinal fat pads with a trend of increased liver weight (**Figure 3a**). The gross appearance of white fats, brown fat, and liver of miR130a-deficient mice and controls were showed in **Figure 3b**. H&E stain of the perigonadal fat (**Figure 3c**) and inguinal fat (**Figure 3d**) showed hypertrophic adipocytes. The adipocyte sizes were larger in the perigonadal fat (**Figure 3e**) and inguinal fat (**Figure 3f**) of miR130a-deficient mice than controls.

**Figure 3.**
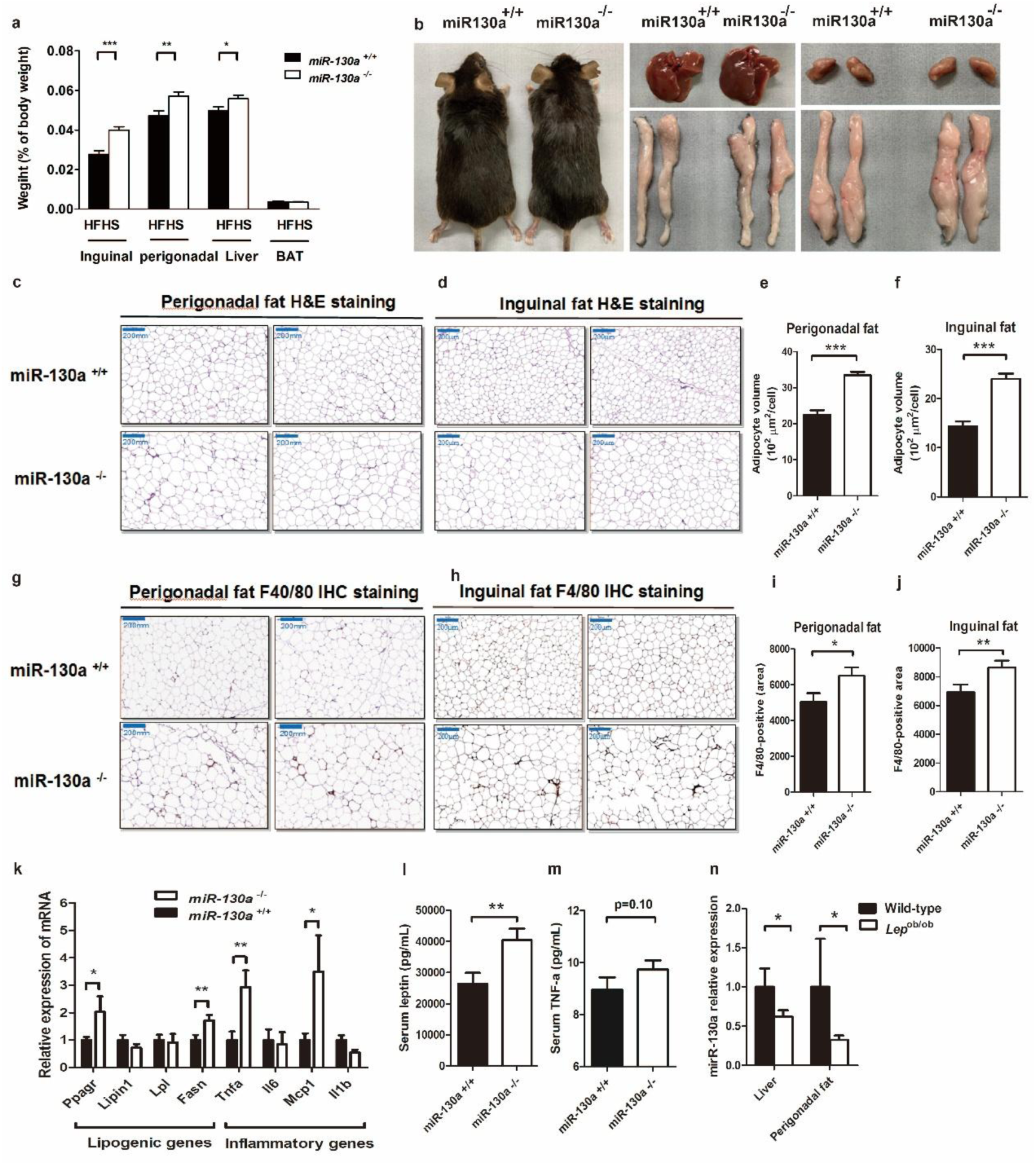
(a)Tissue weight of inguinal fat, perigonadal fat, liver, and brown fat (b) representative gross appearance of inguinal fat, perigonadal fat, and liver of miR130a knockout mice (miR-130a ^−/−^) and controls(miR-130a ^+/+^). H&E stain of (c) perigonadal fat and (d) inguinal fat of miR130a knockout mice (miR-130a ^−/−^) and controls (miR-130a ^+/+^). Adipocyte size of (e) perigonadal fat and (f) inguinal fat of perigonadal fat of miR130a knockout mice and controls. Immunohistochemical stain of F4/80 of (g) perigonadal fat and (h) inguinal fat of miR130a knockout mice and controls. F4/80-positive area of (i) perigonadal fat and (j) inguinal fat of miR130a knockout mice and controls. (k) RT-qPCR showing lipogenic genes and pro-inflammatory genes of perigonadal fat. Fasting serum levels of (l) leptin and (m) TNF-α of miR130a knockout mice (miR-130a ^−/−^) and controls (miR-130a ^+/+^).

Furthermore, immununohistochemical (IHC) stain showed the F4/80-positive infiltrating macrophages in the perigonadal (**Figure 3g**) and inguinal fat (**Figure 3h**) of miR130a-deficient mice are significantly increased with more crown-like necrosis of fat cells than controls (**Figure 3i-3j**). RT-qPCR showed that lipogenic genes including *Pparg* and *Fasn* and pro-inflammatory genes including *Tnfa* and *Mcp1* are increased in the perigonadal fat of miR130a-deficient mice. Consistently, miR130a-deficient mice displayed significantly higher serum letptin levels (**Figure 3l**) and there is a trend of higher serum TNF-α levels than controls (*P*=0.10) (**Figure 3m**). However, serum monocyte chemoattractant protein-1 (MCP-1) levels were not different between mutant and control mice (data not shown).

Oil Red O stain of liver showed that the extent of hepatic steatosis is more severe (**Figure 4a**) and hepatic triglycerides content were higher in miR130a-deficient mice than controls (**Figure 4b**). There is also a trend of higher triglycerides content in quadriceps skeletal muscle from miR130a-deficient mice (P=0.06) than those from controls (**Figure 4c**), suggesting ectopic fat accumulation in skeletal muscle. RT-qPCR showed that lipogenic genes including *Pparg* and *Lpin1* and pro-inflammatory genes including *Tnfa* and *Mcp1* in liver of miR130a-deficient mice (**Figure 4d**). Similarly, Immununohistochemical stain showed the F4/80-positive infiltrating macrophages in the liver were increased in the miR130a-deficient mice than controls (**Figure 4e**). Furthermore, we found that miR130a expression in markedly reduced in the liver and perigonadal fat of the obese *Lep*^ob/ob^ mice compared to wild-type controls (**Figure 4f**).

**Figure 4.**
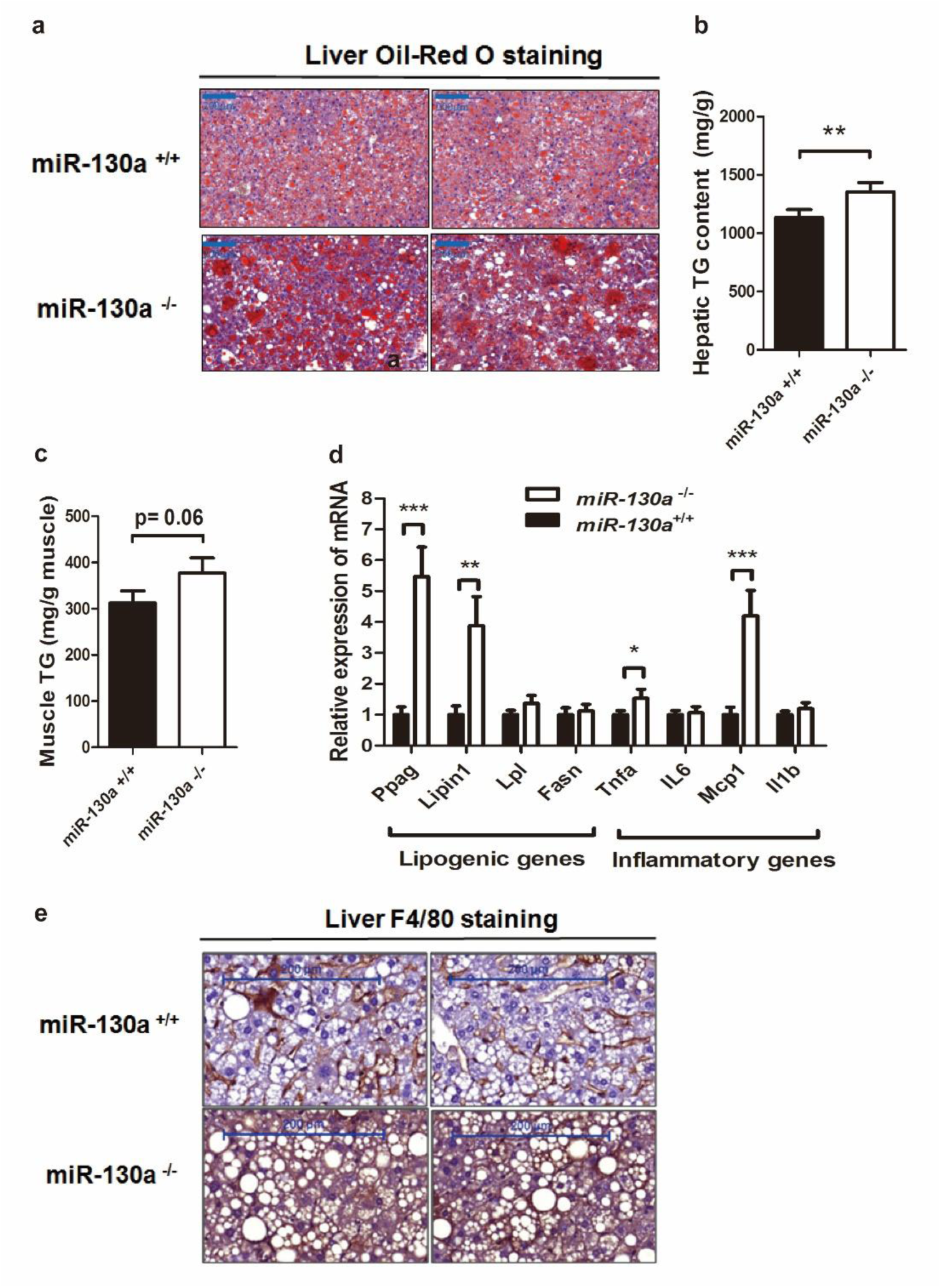
(a)Representative Oil-Red O stain and (b) hepatic triglycerides content of the liver of miR130a knockout mice (miR-130a ^−/−^) and controls (miR-130a ^+/+^) (c) Triglycerides content of the quadriceps muscle of miR130a knockout mice and controls. (d) RT-qPCR showing lipogenic genes and pro-inflammatory genes of the liver of miR130a knockout mice (miR-130a ^−/−^) and controls (miR-130a ^+/+^). (e) Immunohistochemical stains showing the F4/80-positive cells in the liver of miR130a knockout mice (miR-130a ^−/−^) and controls (miR-130a ^+/+^) mice

Importantly, glucose tolerance tests showed and insulin tolerance tests that miR130a-deficient mice are more glucose intolerant (**Figure 5a**) and more insulin resistant (**Figure 5b**) than controls. Immunoblots showed that phospho-akt level are decreased in the perigonadal fat, liver, in skeletal muscle of miR130a-deficient mice after intraperitoneal insulin injection (**Figure 5c**), indicating impaired insulin signaling. In concordance, serum levels of adiponectin, an insulin-sensitizing adipokine were also significantly lower in miR130a-deficient mice than controls (**Figure 5d**). During oral glucose tolerance test, the glucose-stimulated insulin secretion is reduced in miR130a-deficient mice than controls (**Figure 5e**). These data indicated worsened glucose and insulin homeostasis in miR130a-deficient mice.

**Figure 5.**
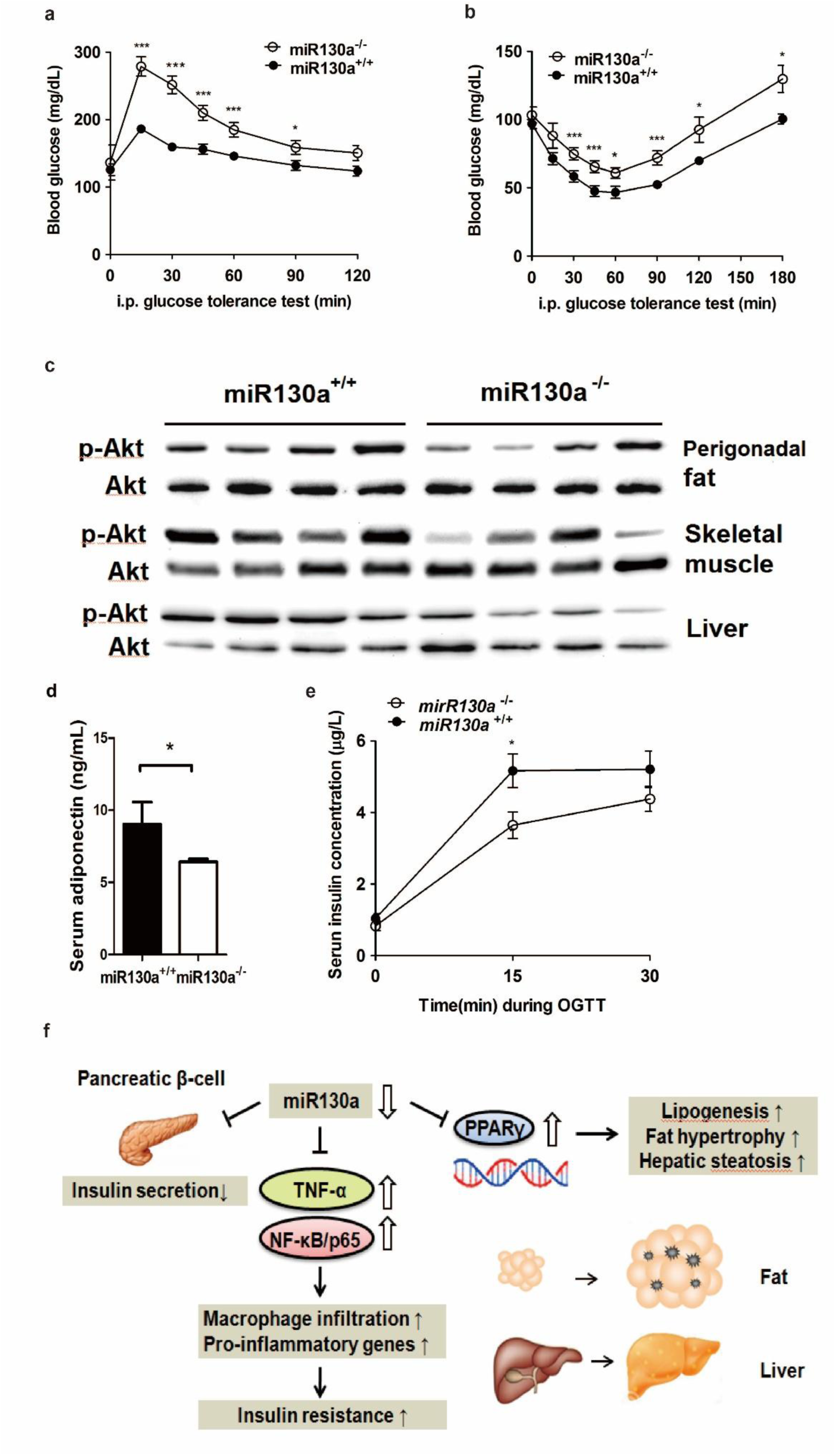
(a)Intraperitoneal glucose tolerance and (b) insulin tolerance tests of miR130a knockout mice (miR-130a ^−/−^) and controls (miR-130a ^+/+^). (c) Phospho-akt expression after insulin injection in the perigonadal fat, skeletal muscle, and liver of miR130a knockout mice (miR-130a ^−/−^) and controls (miR-130a ^+/+^) (d) Fasting serum levels of adiponectin of miR130a knockout mice (miR-130a ^−/−^) and controls (miR-130a ^+/+^).(e) insulin level (o min, 15 min, and 30 min) during oral glucose tolerance test between miR130a knockout mice (miR-130a ^−/−^) and controls (miR-130a ^+/+^). (f) diagram showing the mechanisms by which miR-130a affects systemic fat and glucose metabolism and inflammation.

## Discussions

In this report, we found that miR130a-deficeint mice are prone to develop fat hypertrophy, hepatic steatosis, macrophage infiltration in fat and liver, glucose intolerance, insulin resistance, and impaired glucose-stimulated insulin secretion. These data indicate miR130a exert anti-diabetic effects *in vivo*.

PPARγ is a master regulator of adipogenesis and lipid metabolism. The PPARγ mRNA haw relative short 3’ and 5’ UTRs and therefore few microRNAs have been reported to regulate PPARγ mRNA levels except miR27 and miR20 (21). We and other groups previously reported that miR-130a suppress PPARγ in hepatocytes, preadipocytes, and macrophages by targeting both the coding and the 3’UTR of mRNA (6–11, 13). Over-expression of miR130a suppressed adipocyte differentiation, while inhibition of miR130a enhanced adipogensis and lipognenic gene expression *in vitro* (7). Consistently, here we observed miR130a-deficient mice have hypertropic fat pads, increased lipogenic genes expression, and increased serum leptin concentrations compared to controls. Similarly, we found miR130a-deficient mice had more severe fatty liver with higher hepatic triglycerides content and elevated lipogenic genes expression than controls. There data support that deficiency of miR130a promote lipogenesis *in vivo*. In concordance with these results, we found that miR130a expression in adipose tissue and liver were significantly reduced in the obese *Lep ^ob/ob^* mice compared to controls.

Furthermore, miR130a have been shown to suppress TNF-α mRNA expression in chondrocytes (15) and podocytes (18). TNF-α expression in highly increased while miR130a expression is reduced in tissue of osteoarthritis patients and there is a reverse relationship between TNF-α and miR130a expression in the tissues (15). Mir130a has also been shown to bind to the 3’ UTR of NF-κB/p65, decrease its mRNA stability, and protein levels in macrophages (13). Transfection of miR130a in murine primary hepatocytes stimulated by lipopolysaccharides or free fatty acids greatly inhibits TNF-α expression (13). Here we observed increased macrophage infiltration in the liver and white fat pads of miR130a-deficient mice with increased pro-inflammatory gene expression and there was a trend of elevated serum TNF-α level, indicating miR130a may suppress proinflammatory effects.

Interestingly, previously studies shown that NF-κB drastically stimulated the expression of miR130a in hepatocytes (6) and TNF-α treatment increased miR130a in adipocytes (22). These data suggest a possible negative feedback between miR130a and NF-κB/TNF-α.

Importantly, we found that miR130a-deficient mice exhibited worsen glucose tolerance and insulin sensitivity. This result is consistent with a previous research showing that adenoviral over-expression of miR130a improved glucose tolerance and insulin sensitivity in C57BL6/J mice and ameliorated hepatic steatosis in *Lep ^db/db^*mice (23). However, it is counterintuitive that despite we observed increased expression of PPARγ expression in miR130a-deficient mice, systemic insulin resistance is increased but not improved. This is reminiscent of two independent seminal studies showing that PPARγ heterozygous knockout mice have improved sensitivity than wild-type mice (24, 25). The mechanism by which PPARγ haploinsufficiency improved insulin sensitivity is attributed to decreased fat mass and reduced ectopic triglycerides in skeletal muscle, which alleviates insulin resistance (26). In our study, miR130a-deficient mice displayed increased lipogenesis in white fat and liver with ectopic triglycerides deposition in skeletal muscle owing to increased PPARγ expression. This may explain, at least in part, that miR130a-deficient mice developed insulin resistance despite of increased PPARγ expression. Additionally, deletion of miR130a promotes inflammatory response, which is critical for the development of insulin resistance. The reduced insulin-secretion caused by miR130a deficiency additionally worsened glucose tolerance. The multiple targets of miR130a all together lead to the diabetic phenotypes of miR130a-deficient mice. In conclusion, our results show and an anti-diabetic effect of mirR130a *in vivo*.

## Supporting information

Appedix data

## Notes

### Competing Interest Statement

The authors have declared no competing interest.

