## Supplementary material for "Deficiency of miR130a leads to fat hypertrophy, hepatic steatosis, insulin resistance and glucose intolerance in mice": Appedix data

### Supplementary Figures

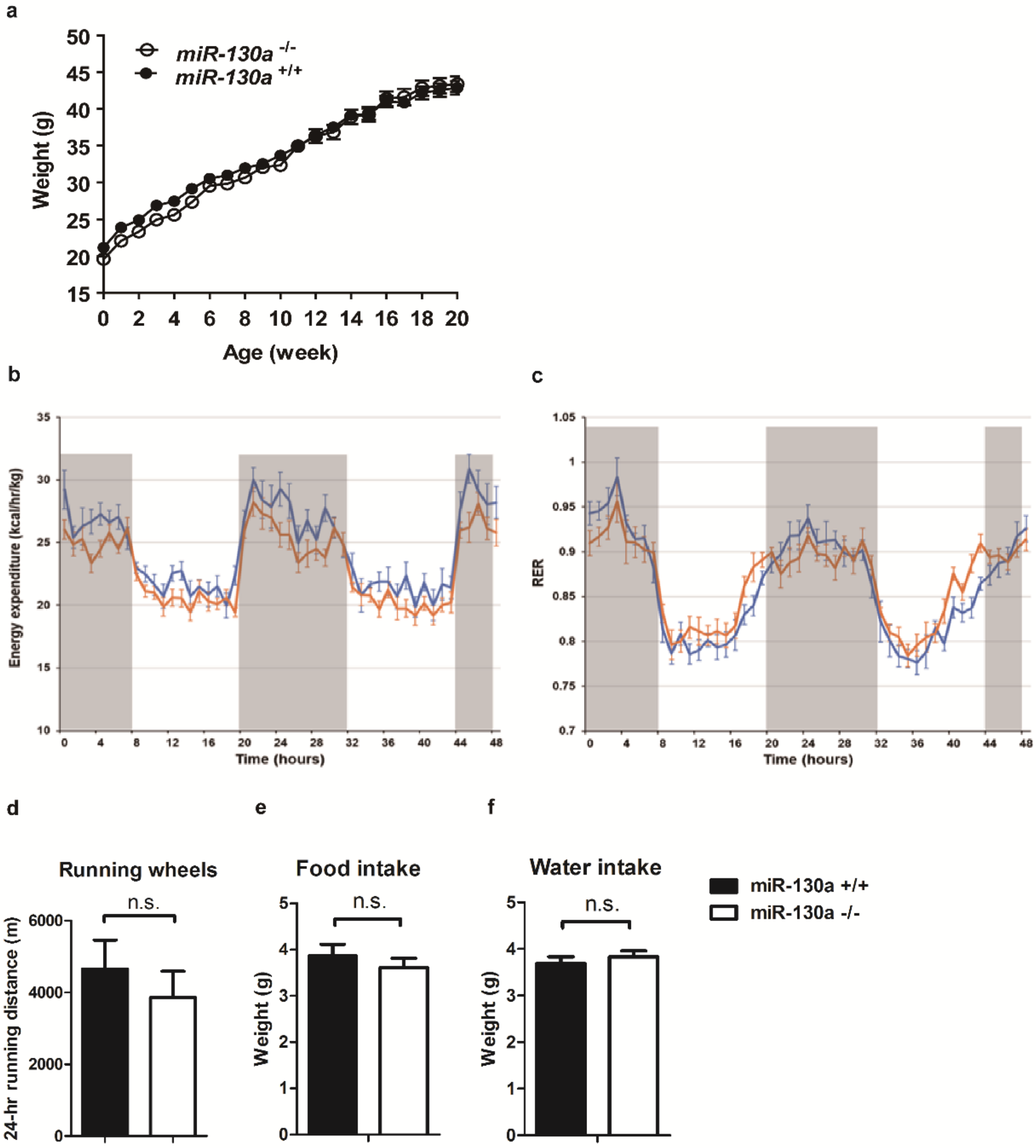


**Supplementary Figures.** (a)Body weight of miR130a knockout mice (miR130a $^{-/-}$) and controls (miR130a $^{+/+}$) on high-fat high-sucrose diet (HFHSD) (b) Energy expenditure and (c) respiratory quotient (RER) of miR130a knockout mice (miR130a $^{-/-}$) and controls (miR130a $^{+/+}$) on HFHSD measure by indirect calorimetry (d) food and water intake and (e) physical activity measured by running wheel rotations of miR130a knockout mice (miR130a $^{-/-}$) and controls (miR130a $^{+/+}$).

### Supplementary Table 1

The primers used in RT-qPCR

| Mouse Gene | 5'-Forward sequence-3' | 5'-Reverse sequence-3' |
|---|---|---|
| *Pparg* | CAAGAATACCAAAGTGCGATCAA | GAGCTGGGTCTTTTCAGAATAATAAG |
| *Lipin1* | CGAGCCACCTCTCTCTCTTGT | GCCCTGGTACTGGGTAGTGAC |
| *Lpl* | TTCCAGCCAGGATGCAACA | GGTCCACGTCTCCGAGTCC |
| *Fasn* | CCCTTGATGAAGAGGGATCA | GAACAAGGCGTTAGGGTTGA |
| *Tnfa* | TACTGAACTTCGGGGTGATTGGTCC | CAGCCTTGTCCCTTGAAGAGAACC |
| *Il6* | TAGTCCTTCCTACCCCAATTTCC | TTGGTCCTTAGCCACTCCTTC |
| *Mcp1* | GCTACAAGAGGATCACCAGCAG | GTCTGGACCCATTCCTTCTTGG |
| *Il1b* | CACAGCAGCACATCAACAAG | GTGCTCATGTCCTCATCCTG |
